# Decreased neutralization of SARS-CoV-2 global variants by therapeutic anti-spike protein monoclonal antibodies

**DOI:** 10.1101/2021.02.18.431897

**Authors:** Takuya Tada, Belinda M. Dcosta, Hao Zhou, Ada Vaill, Wes Kazmierski, Nathaniel R. Landau

## Abstract

Monoclonal antibodies against the SARS-CoV-2 spike protein, notably, those developed by Regeneron Pharmaceuticals and Eli Lilly and Company have proven to provide protection against severe COVID-19. The emergence of SARS-CoV-2 variants with heavily mutated spike proteins raises the concern that the therapy could become less effective if any of the mutations disrupt epitopes engaged by the antibodies. In this study, we tested monoclonal antibodies REGN10933 and REGN10987 that are used in combination, for their ability to neutralize SARS-CoV-2 variants B.1.1.7, B.1.351, mink cluster 5 and COH.20G/677H. We report that REGN10987 maintains most of its neutralization activity against viruses with B.1.1.7, B.1.351 and mink cluster 5 spike proteins but that REGN10933 has lost activity against B.1.351 and mink cluster 5. The failure of REGN10933 to neutralize B.1.351 is caused by the K417N and E484K mutations in the receptor binding domain; the failure to neutralize the mink cluster 5 spike protein is caused by the Y453F mutation. The REGN10933 and REGN10987 combination was 9.1-fold less potent on B.1.351 and 16.2-fold less potent on mink cluster 5, raising concerns of reduced efficacy in the treatment of patients infected with variant viruses. The results suggest that there is a need to develop additional monoclonal antibodies that are not affected by the current spike protein mutations.

## Introduction

Monoclonal antibody therapies for the treatment of COVID-19 have been found to reduce virus loads and alleviate symptoms when given shortly after diagnosis^1,2^. The REGN-COV2 therapy developed by Regeneron Pharmaceuticals is a two recombinant monoclonal antibody cocktail consisting of REGN10933 and REGN10987^3,4^ while the Eli Lilly therapy is based on a single antibody LY-CoV016^5^. The antibodies bind epitopes within the receptor binding domain (RBD) of the Wuhan-Hu-1 spike protein. The rapid evolution of SARS-CoV-2 variants with mutations in the viral S gene that encodes the spike protein raises concerns that monoclonal antibody therapies could lose effectiveness against viruses for which the spike protein has mutations that alter the amino acid sequences of the epitopes bound by the antibodies.

Following the isolation of Wuhan-Hu1 SARS-CoV-2 in December 2019, the virus has continued to further evolve as it adapts to the human host. A variant with a D614G mutation^6^ the spike protein which was identified in January, 2020 and by May became the predominant strain world-wide with a prevalence of >97%. The amino acid residue, which is located near the S1:S2 processing site, reduces S1 subunit shedding from virions, has increased infectivity and results in higher virus loads^7–9^. Additional variants containing the D614G mutation with increased transmissibility were subsequently identified. The B.1.1.7 lineage (VOC-202012/01) variant identified in patients in the United Kingdom^10–12^ encodes a spike protein with 8 mutations in addition to D614G (Δ69-70, Y144Del, N501Y, A570D, P681H, T716I, S982A and D1118H). N501Y is one of six ACE2 contact residues and has been shown to increase affinity for ACE2^13^ by hydrogen bonding with ACE2 Y41^14^; the Δ69-70 deletion in the N-terminal domain is found in multiple independent lineages^15^; and P681H lies adjacent to the furin cleavage site suggesting a role in spike protein processing. The B.1.351 lineage variant identified in patients in South Africa rapidly became the predominant circulating genotype^16^. The virus encodes a spike protein that is more heavily mutated than B.1.1.7 with 9 mutations (L18F, D80A, D215G, L242-244del, R246I, K417N, E484K, N501Y and A701V) three of which (K417N, E484K and N501Y) are in the RBD. E484K, like N501Y, lies in the receptor binding motif (RBM) that directly contacts specific ACE2 residues. K417N, while not contributing to ACE2 binding, is an epitope for neutralizing antibodies, as is E484K, and thus may have been selected for evasion of the humoral response^17–21^. Based on phylogenetic tree branch-length, it has been suggested that the variant arose through the prolonged virus replication in an immunocompromised individual^15^. Additional variants found to be circulating in the human population include the European isolate 20A.EU2^22^, Columbus, Ohio variant COH.20G/677H and the mink cluster 5 variant found in domesticated minks in Denmark with the potential for transfer into humans^23^.

Recent findings have demonstrated partial escape of the B.1.351 variant, and to a lesser extent, B.1.1.7, from neutralization by the serum antibodies of convalescent patients and by antibodies elicited by the Pfizer-BioNtech BNT162b2 and Moderna mRNA-1273 mRNA vaccines that encode trimerized spike proteins^24 25^. The decreased neutralizing titers against B.1.351 were largely the result of the E484K mutation, an amino acid residue that serves as a contact point for ACE2^26–28^.

In this study, we analyzed neutralizing titers of REGN10933 and REGN10987 for viruses with the SARS-CoV-2 variant spike proteins. The results showed that REGN10933 maintains neutralizing activity against B.1.1.7 but has lost neutralizing activity against virus with the B.1.351 and mink cluster 5 spike proteins. Analysis of viruses with the individual B.1.351 mutations mapped the escape to E484K and K417N, residues that lie within the RBD. REGN10987 maintains most of its neutralizing activity against virus with the B.1.1.7, B.1.351 and mink cluster 5 variants, although a small but significant decrease in neutralizing titer was noted against B.1.351 and mink cluster 5 spike proteins. As a result of the decreased activity of both antibodies, the combination of REGN10933 and REGN10987 was decreased in neutralizing titer by 9.1-fold against B.1.351 and 16.2-fold against mink cluster 5.

## Results

The increasing prevalence of highly transmissible variants with mutations in the spike protein RBD raises concerns that the therapy could become less effective should any of the mutations lie within the epitopes targeted by the monoclonal antibodies. To address this question, we tested the neutralizing activity of REGN10933 and REGN10987 on viruses with the variant spike proteins. Neutralizing activity was measured with lentiviral virions pseudotyped with the variant spike proteins, an approach that allows for accurate measurement of neutralizing titers without the need for BSL-3 containment and provides a means to rapidly generate viruses with novel spike proteins variants. Neutralizing titers measured with lentiviral pseudotyped viruses closely match those determined in live virus assays as was shown in a comparative analysis of over 100 convalescent sera analyzed in parallel by both approaches^29^. In this study, we used lentiviral pseudotypes with the parental D614G, B.1.1.7, B.1.351, mink cluster 5 and COH.20G/677H spike proteins and viruses with each of the individual component point mutations and deletions (**Supplementary Figure. 1**).

Analysis of the neutralizing activity of REGN10987 showed that it neutralized D614G with an IC_50_ of 19.4 ng/ml (**Figure. 1A** **and** **Table. 1**). It neutralized B.1.1.7 (Δ69-70-N501Y-P681H) and COH.20G/677H with a similar titers and neutralized B.1.351 and mink cluster 5 spike proteins with slightly higher IC_50_ (2.2-fold and 2.8-fold, respectively). Viruses with each of the single B.1.1.7 mutations were similarly neutralized as were those of B.1.351 and mink cluster 5. Analysis of REGN10933 showed that it was highly active against D614G, B.1.1.7 and COH.20G/677H with an IC_50_ of 7.4, 8.4 and 6.0, respectively, but had weak activity against B.1.351 and mink cluster 5 with an IC_50_ 76.3-fold and 214.9-fold higher, respectively, than that of D614G (**Figure. 1B** **and** **Table. 1**). Analysis of the single mutations of B.1.351 showed that escape from REGN10933 was due to the K417N and E484K, each of which on its own was sufficient. Analysis of spike proteins with the single mutations of the mink cluster 5 variant showed that the escape was caused by Y453F (**Figure. 1B** **and** **Table. 1**). Analysis of the neutralizing activity against the Columbus Ohio variant COH.20G/677H showed a titer comparable to D614G (**Figure. 1** **and** **Table. 1**).

**Figure.1.**
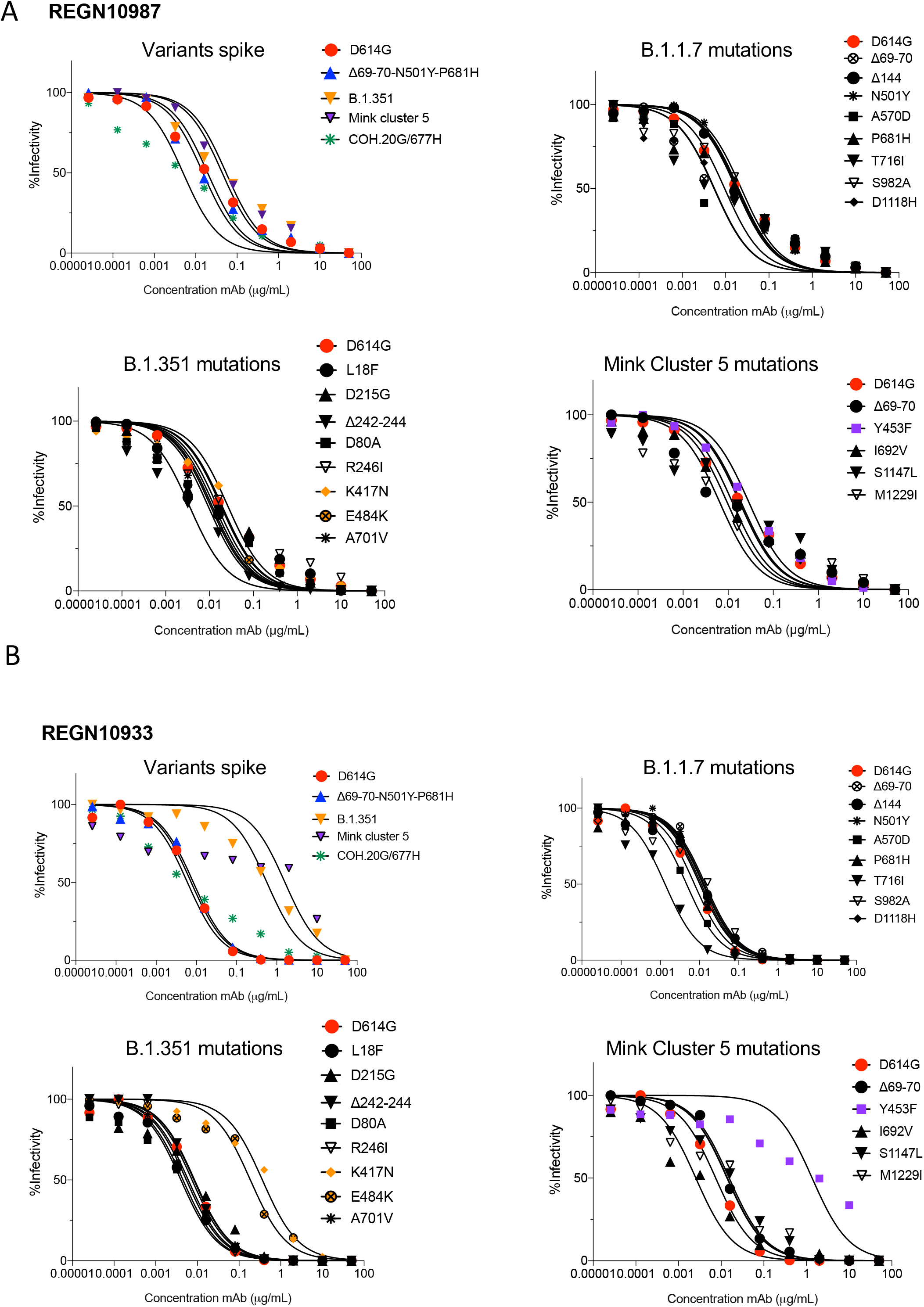
Neutralization of viruses with variant spike proteins by REGN10933 and REGN10987. Neutralization of lentiviral pseudotypes with B.1.1.7, B.1.351 and mink cluster 5 or D614G spike protein by REGN10933 and REGN10987 was measured in ACE2.293T cells. (A) Neutralization curves of viruses with B.1.1.7 (Δ69-70-N501Y-P681H), B.1.351, Mink Cluster 5 and COH.20G/677H (top left), individual B.1.1.7 mutations (top right), individual B.1.351 (bottom left) and Mink Cluster 5 mutations (bottom right) by REGN10987. (B) Neutralization curves of viruses with B.1.1.7 (Δ69-70-N501Y-P681H), B.1.351, Mink Cluster 5 and COH.20G/677H (top left), individual B.1.1.7 mutations (top right), individual B.1.351 (bottom left) and Mink Cluster 5 mutations (bottom right) by REGN10933. The experiment was repeated twice with similar results.

**Table.1.**

|  |  | REGN10933 | REGN10987 | REGN10933+<br>REGN10987 |
| --- | --- | --- | --- | --- |
|  |  | IC50 (ng/mL) |  |  |
| **D614G** |  | 7.44 | 19.42 | 1.69 |
| **B.1.1.7** | **Δ69-70-N501Y-P681H** | 8.40 | 14.94 | 2.18 |
|  | Δ69-70 | 14.03 | 9.49 |  |
|  | Δ144 | 11.11 | 19.79 |  |
|  | N501Y | 12.22 | 18.40 |  |
|  | A570D | 4.68 | 5.52 |  |
|  | P681H | 10.16 | 17.47 |  |
|  | T716I | 1.32 | 5.41 |  |
|  | S982A | 12.92 | 23.00 |  |
|  | D1118H | 10.17 | 17.56 |  |
| **B.1.351** | **B1.351** | 567.60 | 43.74 | 15.45 |
|  | L18F | 4.51 | 9.28 |  |
|  | D80A | 4.02 | 11.46 |  |
|  | D215G | 8.56 | 12.85 |  |
|  | Δ242-244 | 8.26 | 3.48 |  |
|  | R246I | 6.04 | 18.27 |  |
|  | K417N | 362.6 | 26.17 |  |
|  | E484K | 54.29 | 10.55 |  |
|  | N501Y | 12.22 | 18.450 |  |
|  | A701V | 5.43 | 14.98 |  |
| **Mink cluster 5** | **Mink Cluster 5** | 1599 | 54.33 | 27.44 |
|  | Δ69-70 | 14.03 | 9.49 |  |
|  | Y453F | 1355.0 | 28.81 |  |
|  | I692V | 2.54 | 12.40 |  |
|  | S1147L | 15.44 | 17.99 |  |
|  | M1229I | 13.72 | 6.49 |  |
| **COH.20G/677H** |  | 5.98 | 4.96 |  |
| **F486S** |  | >2000 | 8.49 | 8.05 |

Possible escape mutants

In light of the escape of B.1.351 from neutralization by REGN10933, we tested the neutralizing activity of the combination of REGN10933 and REGN10987 which constitutes the REGN-COV2 cocktail on viruses with the variant spike proteins (**Figure. 2** **and** **Table. 1**). The combination of REGN10933 and REGN10987 was highly potent against D614G with an IC_50_ of 1.69 ng/ml, and appeared to be slightly synergistic as the neutralizing titer was higher than of each antibody alone. Neutralizing titers for the mixture against B.1.351 and mink cluster 5 were reduced 9.14- and 16.2-fold compared to D614G, respectively, a result that reflects the large decrease in neutralizing titer for REGN10933 on both variants combined with a minor decrease in neutralizing titer by REGN10987 on both variants (**Figure. 2** **and** **Table. 1**). Analysis of the single point mutations showed that the reduction in neutralizing titer was caused by both E484K and Y453F mutations.

**Figure. 2.**
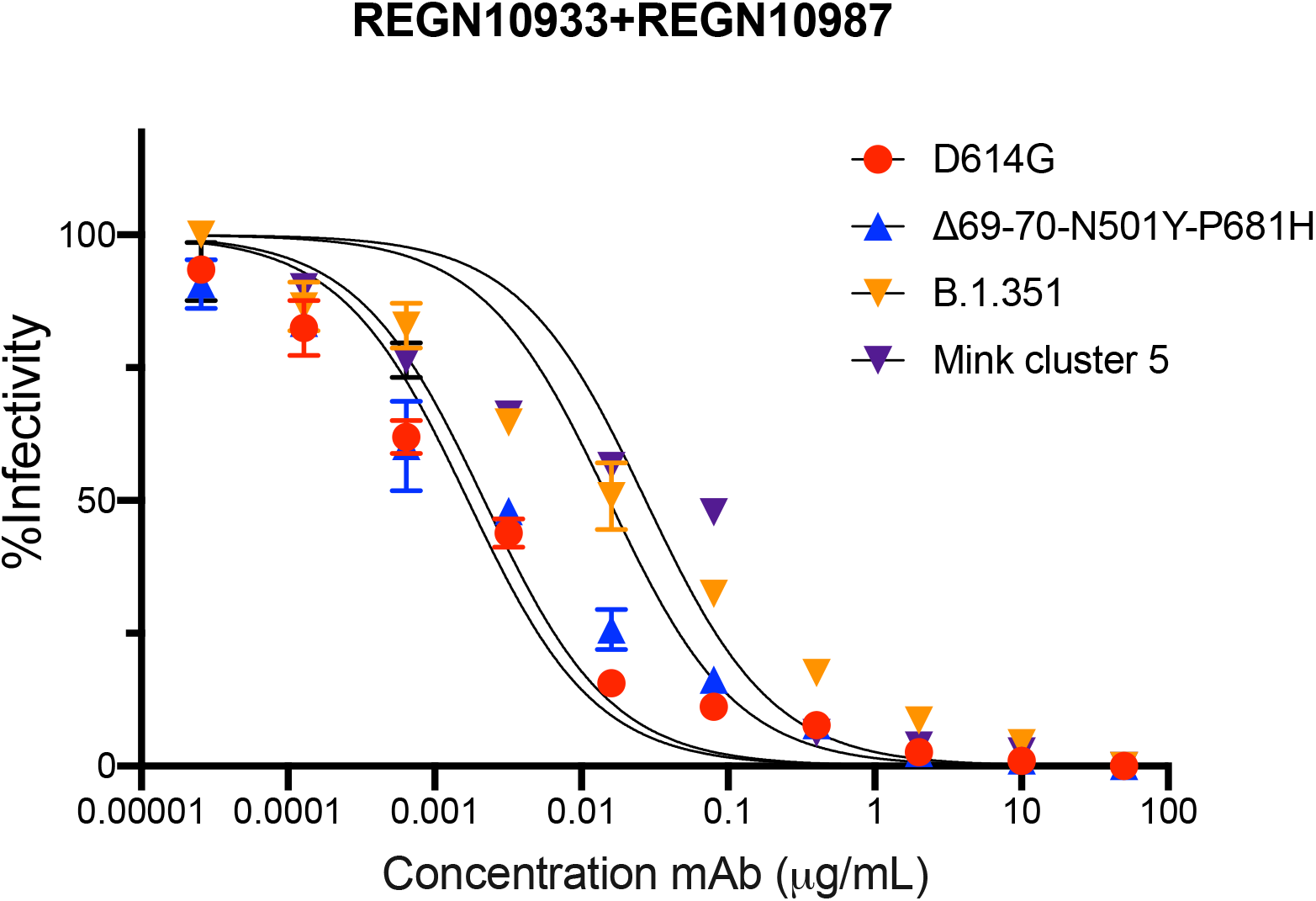
Neutralization of viruses with B.1.1.7, B.1.351 and mink cluster 5 variant spike proteins by the REGN10933 and REGN10987 cocktail. Neutralization of viruses with B.1.1.7, B.1.351 and mink cluster 5 or by D614G spike proteins by a 1:1 mixture of REGN10933 and REGN10987 was measured. The experiment was repeated twice with similar results.

## Discussion

We report here that REGN10933, one of the two monoclonal antibodies that constitutes the REGN-COV2 therapy for SARS-CoV-2, has lost neutralizing activity against viruses with the South Africa B.1.351 and mink cluster 5 variant spike proteins. The other antibody, REGN10987, maintains most of its neutralization activity against the B.1.351 and mink cluster 5 variants. Analysis of viruses with spike proteins containing the individual B.1.351 spike protein mutations showed that the escape was caused by the K417N and E484K mutations in the RBD, either of which separately prevents neutralization, findings that are consistent with those recently reported by Wang *et al^27^*. As a result of the decreased potency of both antibodies, the combined REGN10933 and REGN10987 cocktail had a decrease in neutralizing titer of 9.1-fold against B.1.351 and 16.2-fold against mink cluster 5.

REGN10933 and REGN10987 bind to non-overlapping sites on the RBD^3^. REGN10933 binds at the top of the RBD, blocking the interaction with ACE2 while REGN10987 binds to the side of the RBD and does not overlap with the ACE2 binding site^3^. The spike protein mutations that affect REGN10933 (E484K, K417N and Y453F) cluster on the side of the RBD to which the antibody binds (**Figure. 3**). In addition, we found that the mutation F486S prevents neutralization by REGN10933 (**Supplementary Figure. 2** **and** **Table. 1**). The amino acid is located close to E484 and has been reported to affect ACE2 binding^30^. The relative sensitivity of REGN10933 to mutations in spike protein variants may result from selective pressure to optimize ACE2 interacting amino acids in the RBD such as E484, a pressure that is not as great for the amino acids bound by REGN10987 which lie on the other face of the RBD.

**Figure. 3.**
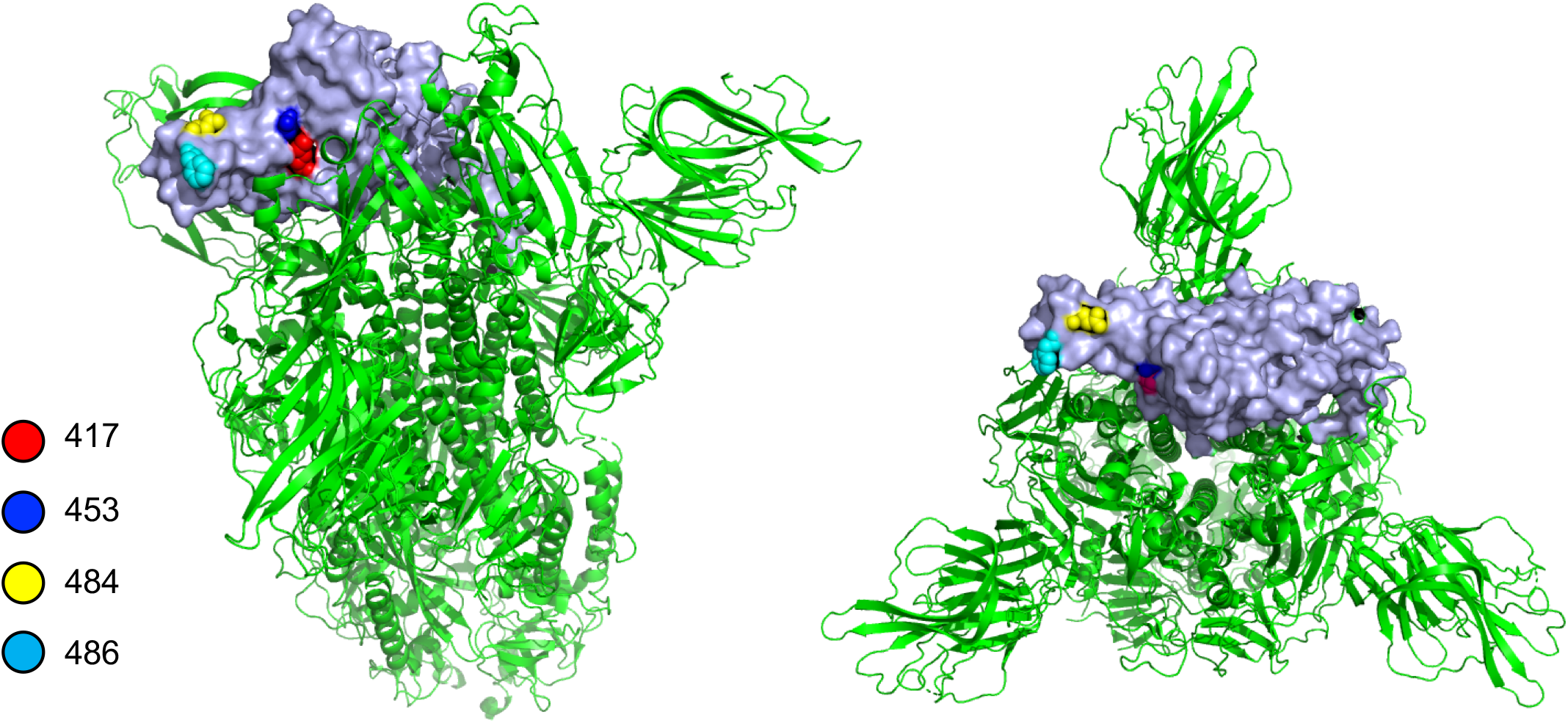
Location of amino acid residue causing escape from REGN10933 in the SARS-CoV-2 spike protein. Side (left) and top (right) views of the prefusion structure of the SARS-CoV-2 spike protein. The location of amino acid residues 417, 453, 484 and 486 in the RBD that cause escape from REGN10933 are shown (a single RBD is shown in grey).

Previous studies have explored binding of REGN10933 and REGN10987 to the RBD and analyzed mutations that allow for escape. Using a deep mutational scanning method, Starr *et al*. found that mutations at residue 486 escaped neutralization by REGN10933 whereas mutations at residues 439 and 444 escaped neutralization by REGN10987^31^. A single mutation, E406W, allowed for escape from both antibodies although, interestingly, the residue is not located within the epitope bound by either antibody. Analysis of spike protein mutations that occurred in a treated immunocompromised patient revealed additional mutations that allowed for escape from either antibody^32^. While these mutations allow for escape, they may not become problematic for therapy as they are not selected by immune pressure and may have unrecognized effects on viral fitness that reduce transmissibility.

It is not clear whether the reduced neutralizing titer of the REGN10933 and REGN10987 cocktail will translate into a loss of effectiveness of REGN-COV2 therapy for individuals infected with the B.1.351 variant. The two antibody cocktail has an IC_50_ of 15.4 ng/ml which is still substantial. The findings highlight the benefit of a two antibody cocktail as therapy with the single REGN10933 would have lost effectiveness for use in patients infected with the B.1.351 variant and would be problematic in populations in which the variant was prevalent. While REGN10987 has so far been unaffected by spike protein mutations, it would be advantageous to develop additional monoclonal antibodies that retain neutralizing activity against current spike protein variants, particularly in light of the increasing prevalence of variants with mutations in critical amino acid positions of the spike protein. The results presented here highlight the importance of continued surveillance for SARS-CoV-2 variants and for testing the sensitivity of variants to anti-spike protein neutralizing antibodies in clinical use as well as their ability to be neutralized by vaccine elicited antibodies. These findings highlight the need to identify antibodies against highly conserved spike protein epitopes which the virus cannot readily mutate.

## Methods

### Plasmids

pLenti.GFP.NLuc dual GFP/nanoluciferase lentiviral vector, pcCOV2.Δ19S codon-optimized SARS-CoV-2 spike gene expression vector based on the Wuhan-Hu-1/2019 amino acid sequence with a termination codon at position 1255, HIV-1 Gag/Pol expression vector pMDL and HIV-1 Rev expression vector pRSV.Rev have been previously described^33^. Point mutations in pcCOV2.Δ19S open reading frame were introduced by overlap extension PCR. All plasmid sequences were confirmed by DNA nucleotide sequence analysis.

### Cells

293T cells were cultured in Dulbecco’s modified Eagle medium (DMEM) supplemented with 10% fetal bovine serum (FBS) and penicillin/streptomycin (P/S) at 37°C in 5% CO_2_. ACE2.293T cells are clonal cell-line that expresses high levels of human ACE2 and have been previously described ^29,33^.

### Monoclonal antibody production

cDNAs encoding REGN10933 and REGN10987 were synthesized using the published sequences of the antibody variable heavy and light chains fused to IgG1 heavy chain and lambda light chain, respectively and cloned into pcDNA3.1 (Invitrogen). The proteins were produced in transfected Freestyle 293 cells and collected from the cell supernatant after four days. The antibodies were purified by on an AKTA prime FPLC with HiTrap Pro A 5cc column. The proteins were tested for purity by SDS-PAGE, quantified by BCA assay and tested for spike protein binding by Bio-layer Interferometry on an Octet Detection System.

### SARS-CoV-2 spike lentiviral pseudotypes

SARS-CoV-2 spike protein pseudotyped lentiviral stocks were produced by cotransfection of 293T cells with pMDL, pLenti.GFP-NLuc, pcCoV2.S-Δ19 (or variants thereof) and pRSV.Rev as previously described^33^. Virus stocks were normalized for reverse transcriptase activity^34^. Pseudotyped virus infections were done with 1 × 10^4^ cells/well in 96 well tissue culture dishes at an MOI=0.2^33^. Luciferase activity was measured after 2 days using Nano-Glo luciferase substrate (Promega) and plates were read in an Envision 2103 microplate luminometer (PerkinElmer). To measure antibody neutralization, antibodies were serially diluted 5-fold and then incubated for 30 minutes at room temperature with pseudotyped virus (corresponding to approximately 2.5 × 10^7^ cps luciferase) in a volume of 50 μl. The mixture was added to 1 × 10^4^ ACE2.293T cells (corresponding to an MOI of 0.2) in a volume of 50 μl in a 96 well culture dish. After 2 days, the medium was removed and Nano-Glo luciferase substrate (Nanolight) was added to wells. Luminescence was read in an Envision 2103 microplate luminometer (PerkinElmer).

### Quantification and Statistical Analysis

All experiments were performed in technical duplicates or triplicates and data were analyzed using GraphPad Prism (Version 8). The PDB file of D614G SARS-CoV-2 spike protein (7BNM) was downloaded from the Protein Data Bank. 3D view of protein was obtained using PyMOL.

## Acknowledgements

The work was funded by grants from the NIH to N.R.L. (DA046100, AI122390 and AI120898). T.T. was supported by the Vilcek/Goldfarb Fellowship Endowment Fund.

## Author contributions

T.T. and N.R.L. conceived and designed the project. T.T., B.M.D and H.Z. carried out the experiments and analyzed the data. A.V., W.K. provided the monoclonal antibodies REGN10933 and REGN10987. T.T. and N.R.L wrote the manuscript. All authors provided critical comments on manuscript.

---

**Supplementary Figure 1.**
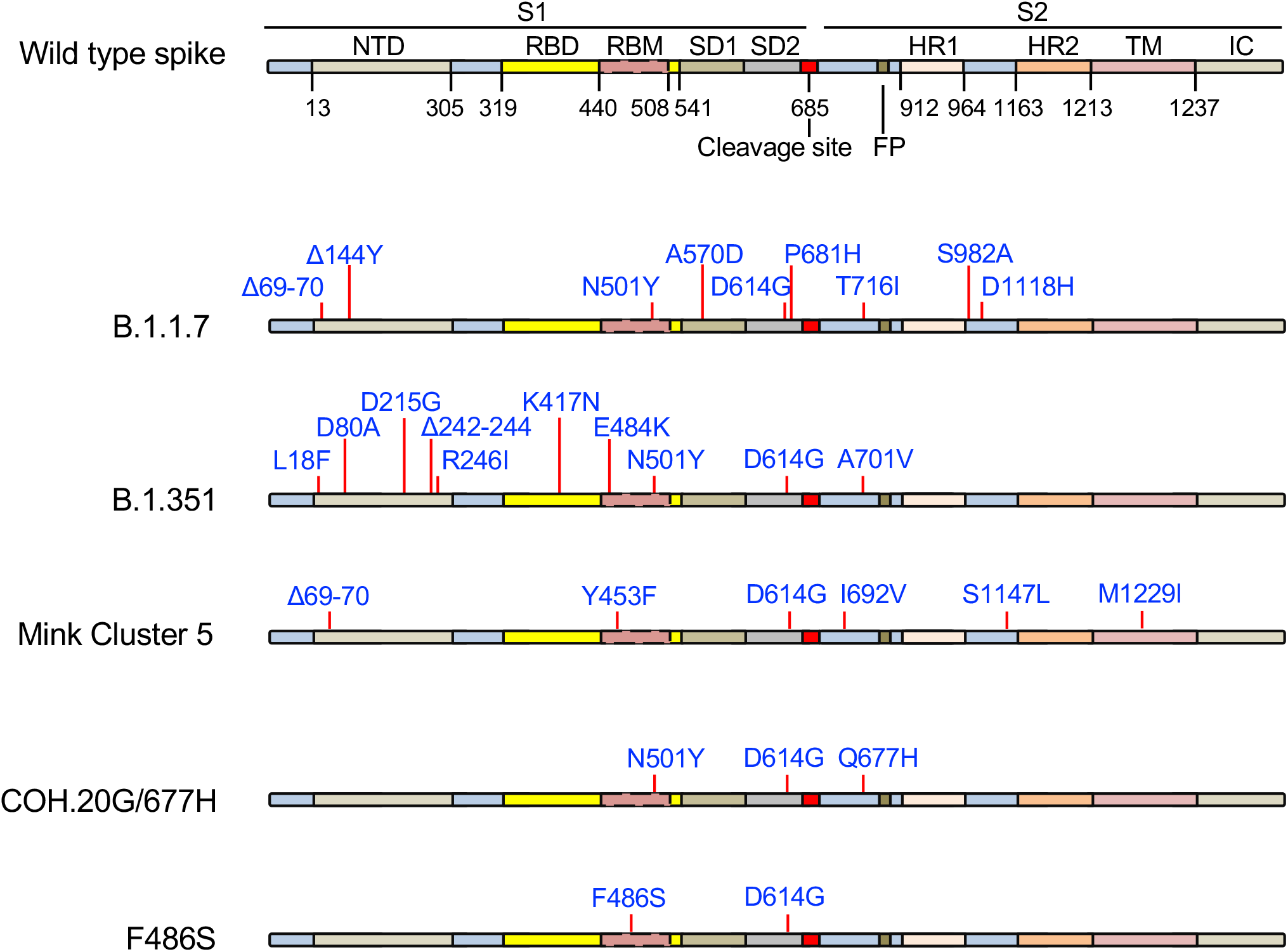
The structure of the SARS-CoV-2 spike protein and location of mutated amino acid residues of the variant spike proteins is diagrammed. The domains of the spike protein are indicated as NTD, N-terminal domain; RBD, receptor-binding domain; RBM, receptor-binding motif; SD1 subdomain 1; SD2, subdomain 2; FP, fusion peptide; HR1, heptad repeat 1; HR2, heptad repeat 2; TM, transmembrane region; IC, intracellular domain.

**Supplementary Figure 2.**
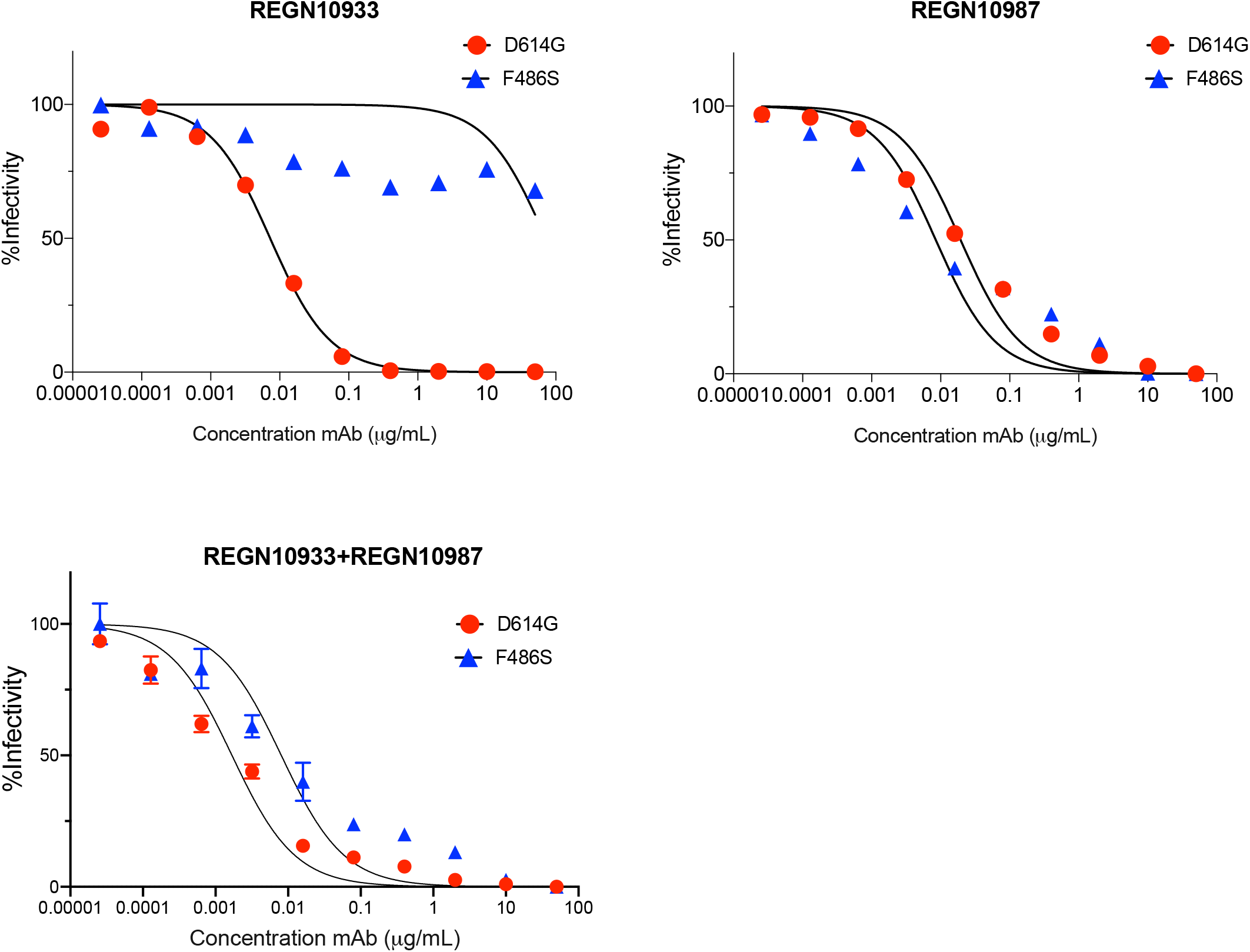
Neutralization curves for REGN10933 and REGN10987 on virus with F486S mutated spike protein. Neutralization by REGN10933 and REGN10987 of lentiviral pseudotyped virions with F486S or D614G mutations in the spike protein were analyzed. F486S is not one of the major circulating variants.

